# Gabapentinoids promote striatal dopamine release and rescue multiple deficits of a mouse model of early Parkinson’s

**DOI:** 10.1101/2025.09.16.675586

**Authors:** Katherine R. Brimblecombe, Adam Loyd Harris, Nora Bengoa-Vergniory, Bethan O’Connor, Lucille Duquenoy, Rishi Anand, Emanuel F. Lopes, Bradley M. Roberts, Lauren Burgeno, Mark E. Walton, Stephanie J. Cragg

## Abstract

Neuronal entry and handling of intracellular calcium have long-been hypothesised to burden vulnerable dopamine neurons in Parkinson’s disease. However, no treatments for Parkinson’s target calcium biology. Gabapentinoid drugs bind to α_2_δ subunits of voltage-gated calcium channels (VGCCs) and are licensed for neurological disorders including dopamine-dysregulated restless leg syndrome, suggesting their potential utility to modify both calcium biology and dopamine signalling. We therefore tested whether gabapentinoids modulate dopamine signalling, underlying VGCC-dependence, and potential for treating Parkinson’s. In mouse striatum, we reveal that gabapentinoids *ex vivo* promote dopamine release, via sex-specific dependence on α_2_δ1/2 subunits and alterations to the calcium- and VGCC-subtype-dependence of dopamine release or its tonic inhibition by striatal GABA. *In vivo* administration of gabapentinoids to a mouse model of early Parkinson’s rescued deficits in dopamine release, dysregulation of GABAergic inhibition and dopamine content, and abolished parkinsonian deficits in movement transitions. Thus, gabapentinoids urgently deserve attention for repurposing for Parkinson’s disease.

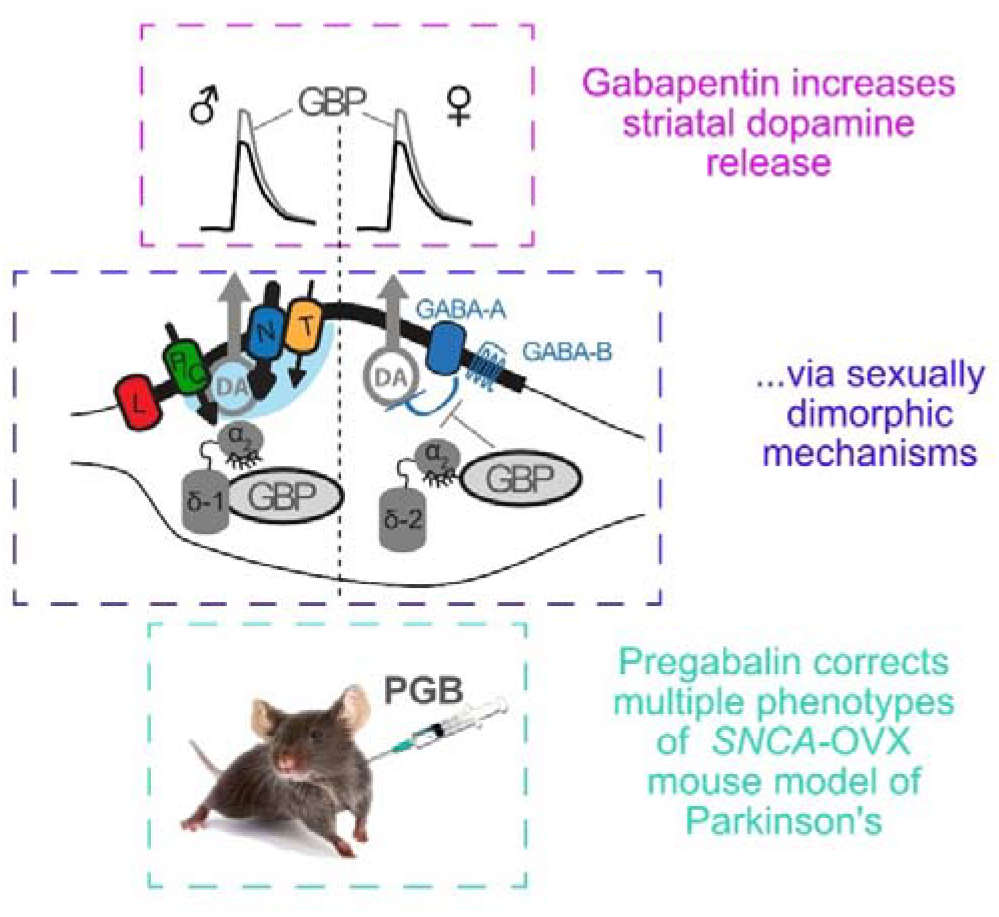

## Introduction

Metabolic stress arising from high intracellular calcium (Ca^2+^) load and weak Ca^2+^-buffering is suggested to be a contributing factor in the loss of selective subpopulations of dopamine (DA) neurons of the substantia nigra pars compacta (SNc) in Parkinson’s disease ^1^. Multiple subtypes of voltage-gated Ca^2+^ channels (VGCCs) have been suggested to contribute to the somatic Ca^2+^ burden of SNc neurons, particularly L-, T- and R-type channels that support somatodendritic subthreshold conductances ^2–4^. Furthermore, a broader range of VGCC types operate to support DA release from striatal axons arising from SNc than from adjacent ventral tegmental area (VTA) neurons ^5^. VGCCs are multi-subunit complexes, consisting of a pore-forming α_1_ subunit, which confers characteristic properties of N-, P/Q, R-, L- and T-subtypes, associated with β, γ and α_2_δ subunits ^6^. The a δ subunits regulate VGCC trafficking, localization and levels in the plasma membrane, and given the high level of membrane turnover in DA axons ^7^, α_2_δ subunits here are likely to be actively governing how VGCCs support DA release. Midbrain DA neurons express α_2_δ1-3 subunits, with at least α_2_δ3 subunit showing relative enrichment in SNc versus VTA neurons ^8,9^. But whether α_2_δ subunits support function of VGCCs in gating DA transmission has not previously been explored.

Gabapentinoid drugs e.g. gabapentin (GBP) and pregabalin (PGB) bind to the RRR-motif contained in α_2_δ-1 and α_2_δ-2 subunits and limit the forward trafficking of the α_1_ subunit. Gabapentinoids can therefore decrease presynaptic VGCC density and Ca^2+^ influx, decreasing release of some neurotransmitters ^10–12^. Gabapentinoids are approved for treatment of multiple disorders including epilepsy, neuropathic pain, sleep disturbances, anxiety and restless leg syndrome, which is associated with DA dysregulation ^13,14^. The clinical efficacy of gabapentinoids is due to their interaction with α_2_δ1/2, which act as subunits of VGCCs. Intriguingly, there is evidence from small scale trials that gabapentinoids might have some clinical efficacy in Parkinson’s patients, including improving clinical UPDRS scores as well as other co-morbidities such as pain ^15–17^, but whether gabapentinoids have a relevant impact on DA transmission, or on underlying α_2_δ regulation of VGCC function in DA axons, that might provide a biological underpinning for further testing, are fundamental and important questions that have not previously been addressed.

Here, we probed whether gabapentinoids impact on DA release and on movement impairments in a mouse model of Parkinson’s. We found that *ex vivo* or *in vivo* administration of gabapentinoids increased evoked DA release in the dorsolateral striatum, involving α_2_δ subunits and distinct mechanisms in male versus female mice. Moreover, in a humanised α-synuclein-overexpressing mouse model of Parkinson’s, *in vivo* pregabalin administration ameliorated locomotor alterations and associated striatal circuit disturbances. These data provide a mechanistic rationale to underscore the potential utility of gabapentinoids for the treatment of Parkinson’s disease that now warrants significant follow-up.

## Materials and Methods

### Animals

All procedures were performed in accordance with the Animals in Scientific Procedures Act 1986 (Amended 2012) with ethical approval from the University of Oxford, and under authority of a Project Licence granted by the UK Home Office. Adult mice (8-20 weeks) of both sexes wild-type C57bl6/J (Charles River, RRID:IMSR_JAX:000664); Homozygous male DATIREScre mice (https://www.jax.org/strain/006660, RRID:IMSR_JAX:006660) were crossed with homozygous Ai95(RL-GCamp6f)-D female mice to generate mice heterozygous for both alleles for Ca^2+^ photometry experiments (age 8-12 weeks of age). *SNCA*-OVX and *Snca*-null mice were generated in house as previously published (Janezic et al., 2013) (OVX https://www.jax.org/strain/023837, RRID:IMSR_JAX:023837; null https://www.jax.org/strain/003692, RRID:IMSR_JAX:003692). *Cacna2d1*-RA and *Cacna2d2*-RA mutants were sourced from Pfizer via Charles River under material transfer agreement ^19,20^ and homozygous adult mice (8-20 weeks) of both sexes were used as experimental animals.

### Slice preparation

Mice were killed by cervical dislocation, the brains removed, and 300 µm coronal striatal slices prepared using vibrating microtome (Leica VT1200 S). For voltammetry slices were prepared in ice-cold HEPES-based buffer saturated with 95% O_2_/5% CO_2_, containing in mM: 120 NaCl, 20 NaHCO_3_, 6.7 HEPES acid, 5 KCl, 3.3 HEPES salt, 2 CaCl_2_, 2 MgSO_4_, 1.2 KH_2_PO_4_, 10 glucose. Slices were incubated at room temperature for ≥ 1 hour in HEPES-based buffer before experiments. For Ca^2+^ photometry, slices were prepared in ice-cold high sucrose cutting solution [(in mM: 194 sucrose, 30 NaCl, 4.5 KCl, 1 MgCl_2_, 26 NaHCO_3_, 1.2 NaH_2_PO_4_, and 10 glucose], and then transferred to aCSF (in mM: 130 NaCl, 2.5 KCl, 26 NaHCO_3_, 1.25 NaH_2_PO_4_, 10 glucose, 2 MgCl_2_-6H_2_O, 2.5 CaCl_2_ and saturated with 95% O_2_ /5% CO_2_) at 32°C for 15 minutes, slices were then returned to room temperature for 45 minutes prior to recording. All procedures were licensed to be carried out at the University of Oxford under the UK Animals (Scientific Procedures) Act 1986.

### Voltammetry

*Ex vivo* DA release was monitored in acute coronal slices using fast-scan cyclic voltammetry (FCV) as previously described ^5^. Slices were superfused in a recording chamber with bicarbonate-buffered artificial cerebrospinal fluid (aCSF) saturated with 95%O_2_/5%CO_2_ at 31–33 °C, containing in mM: 124 NaCl, 26 NaHCO_3_, 3.8 KCl, 0.8-3.6 CaCl_2_ (as stated), 1.3 MgSO_4_, 1.3 KH_2_PO_4_, 10 glucose. To exclude potentially confounding effects of actions of striatal acetylcholine acting at nicotinic receptors on DA release ^49–51^, all data presented here are collected in the presence of nicotinic receptor antagonist DHβE (1 μM). Evoked extracellular DA concentration ([DA]_o_) was monitored by FCV using a Millar voltammeter (Julian Millar, Barts and the London School of Medicine and Dentistry) and single-use carbon-fibre microelectrodes (7-10 μm diameter) fabricated in-house (tip length 50-100 µm). A triangular voltage waveform (range -700 mV to +1300 mV vs. Ag/AgCl) was applied at 800 V/s at a scan frequency of 8 Hz. Electrodes were switched out of circuit between scans. Electrodes were calibrated using 2 μM DA, prepared immediately before calibration using stock solution (2.5 mM in 0.1M HClO_4_ stored at 4 °C). Signals were attributed to DA due to the potentials of their characteristic oxidation (500-600 mV) and reduction (-200 mV) peaks.

### Electrical stimulation

DA recordings were obtained from dorsolateral quadrant of striatum. DA release was evoked by a local bipolar concentric Pt/Ir electrode (25 µm diameter; FHC inc. ME, USA) placed approximately 100 µm from the recording electrode. Stimulus pulses (200 µs duration) were given at 0.6 mA (perimaximal in drug-free control conditions). Stimulations were single pulses (1p) or trains of 5 pulses (5p) at 5-100 Hz as specified and were repeated at 2.5 minute intervals, with 1p stimulations occurring every third stimulation to ensure site stability over time. Each stimulation type was recorded in at least triplicate in each recording site in all experimental conditions. All data were obtained in the presence of the nAChR antagonist, dihydro-β-erythroidine (DHβE, 1 μM) to exclude the powerful modulatory effects of cholinergic interneurons on DA release ^49,50,52,53^.

### Gabapentin incubation

Slices were incubated in either gabapentin-containing aCSF (GBP, 50 μM) or control aCSF for 30 min before being transferred to the recording chamber. The recording electrode was inserted into a non-recording site and an additional 30 min was allowed for the electrode to charge and equilibrate. During this time, the slice was superfused with GBP-containing aCSF (50 μM) or control aCSF. Prior to recording, DHβE (1 μM) was added to the superfusion media. For GBP conditions, GBP (50 μM) was present throughout. The alternative gabapentinoid drug pregabalin (100 µM) was included in the superfusate for 30-45 min prior to recording. The concentration of 50 μM GBP selected was based on previously used concentrations ^22–24^ and can be approximated to 7 μg/mL CSF levels (based on serum concentrations and partition ratios in mice), which is equivalent to a moderate dose of GBP in people of 900 mg/day ^18^. It should be noted that the relationship between [DA]_o_ and stimulation frequency in the absence and presence of GBP was conducted at 1.2 mM [Ca^2+^]_o_ as the relationship between DA release stimulation frequency is enhanced at lower [Ca^2+^]_O_, albeit the relationship remains weak ^5,54^.

Since gabapentinoids act not through acute channel inhibition, but by interfering with VGCC subunit interactions that change channel localization and function, their actions are usually considered to be apparent only after incubation. Relatively acute effects of gabapentinoids are not thought to occur in reduced preparations such as cultured neurons ^21,55^. But the timescale over which GBP acts is likely to vary with experimental conditions. More intact preparations promote α_2_δ function over shorter timescales ^56^, and we based our ∼1 hour GBP incubation time on previous studies which have identified GBP actions in *ex vivo* slice preparations from other brain regions ^22,57^. The function of α_2_δ subunits is thought to be particularly apparent in situations where there is a high degree of endocytosis/membrane turnover ^58^. Notably, a high degree of endocytosis has been shown to occur in DA axons in *ex vivo* striatal slices ^7^ making it likely that GBP actions will be detected over short time scales in this preparation.

### EGTA/BAPTA incubation

Striatal sections were bisected and each hemisphere was incubated for 30 mins at room temperature in aCSF containing 2-hydroypropyl-β-cyclodextrin, 70 µM (Sigma), probenecid, 175 µM, (Sigma), pluronic acid, 0.1% (Life Technologies), and either EGTA-AM, 100 µM (Millipore) or BAPTA-AM, 100 µM (Tocris), or DMSO (vehicle control) ^59,60^ in the absence or presence of GBP (50 μM). Following pre-incubation, hemispheres were incubated for a further 30 minutes in the recording chamber prior to recording. Recordings were alternated between the EGTA-AM/BAPTA-AM-incubated versus non-incubated slice and at paired recording locations. EGTA-AM/BAPTA-AM effects sizes were obtained from peak [DA]_o_ expressed as a percentage of control paired site.

### Ca^2+^ imaging

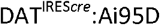 mice expressing GCaMP6f in dopaminergic neurons were used for Ca^2+^ imaging experiments. Following gabapentin or vehicle (aCSF) incubation, striatal slices were hemisected and transferred to a recording chamber at 32 ± 1°C, superfused with aCSF containing in mM: 130 NaCl, 2.5 KCl, 26 NaHCO_3_, 1.25 NaH_2_PO_4_, 10 glucose, 2 MgCl_2_-6H_2_O, 2.5 CaCl_2_ and saturated with 95% O_2_/5% CO_2_. An Olympus BX51WI microscope equipped with a rig a IRIS 9 CMOS camera (Teledyne Photometrics), or a Scientifica Slicescope equipped with a Prime BSI Express sCMOS camera (Teledyne Photometrics), with a 10x/0.3 NA water immersion objective (Olympus), was used to detect changes in GCaMP6f fluorescence resulting from single pulse electrical stimulation in DLS. During acquisition, slices were illuminated using a LED light (λ = 470 nm) at a power of 5 mW (measured under at 10x objective). Images were acquired at 2.5 minute intervals, with a 100 Hz frame rate (10 ms exposure, 2x2 pixel binning), with image acquisition and electrical simulation time-locked using Micro-Manager (2.0) and MC_Stimulus II (Multi Channel Systems) software. Data were analysed using Fiji (version 2.9) and Matlab (version R2020b) to fit a logarithmic curve (a*log(x)+b) to correct for initial decay in fluorescence and to extract fluorescence intensity over time from the region of interest (ROI). F_0_ was defined as the b component of the fitted curve. Data are expressed as ΔF/F_0_.

### Dosing with PGB

Pregabalin (purchased from Sigma) was dissolved in sterile saline in sterile conditions and filtered through 0.22 μm PES membrane (Millipore). Sterile PGB was aliquoted into sterile Eppendorf tubes and stored at -20°C until required. Pairs of *SNCA*-OVX mice were randomly assigned saline or PGB conditions and were acclimatised to handling and scruffing (soft scruff, without mice being lifted) for 3 days and were weighed. Mice were injected sub cutaneously with either saline or PGB 30 mg/kg twice daily for 3 days (day 1: PM; day 2: AM, PM; day 3: AM,PM; day 4: AM) using 29G 1.0 mL insulin needle. Final dose was given 1 hour before experiment conducted (either mouse killed by cervical dislocation for voltammetry experiments or mouse transferred to non-home cage for OFT). Twice daily subcutaneous 30 mg/kg PGB can be estimates as being equivalent to 300-600 mg/day in patients ^61–63^.

### Drugs

BAPTA-AM, dihydro-β-erythroidine (DHβE), GBP, NNC 55-0396, isradipine, ω-Agatoxin IVA and ω-Conotoxin GVIA, CGP-55845 and bicuculline were purchased from Ascent Scientific or Tocris UK; pluronic acid from Life Technologies; EGTA-AM from Millipore. All other reagents were purchased from Sigma Aldrich. Stock solutions were made to 1000-2000x final concentrations in H_2_O (DHβE, GBP, ω-conotoxin GVIA, ω-Agatoxin IVA and NNC 55-0396, cocaine), DMSO (isradipine, EGTA-AM, BAPTA-AM, Bicuculline and CGP-55845) and stored at -20°C. Drugs were diluted to their required concentrations in aCSF immediately prior to use. Drug concentrations were chosen in accordance with previous studies.

### HPLC

After FCV recordings, brain slices were transferred to petri-dish and tissue punches (2 x 1.2mm for NAc) and (2 x 2 mm for DLS) were taken from pairs of SNCA-OVX male mice treated with saline vs PGB (x6) were put in perchloric acid solution (PCA) (200 µL, 0.1 M) and stored at -80 °C. On day of measurement all samples were homogenised using hand-held sonicator and spun at 14,000 G at 4 °C for 15 min. DLS samples were diluted 15X in PCA and NAc samples were run neat and quantified against known concentration standards (100 nM) using 4.6 X 150 mm Microsorb C18 reverse-phase column (Varian) and Decade II ECD with a Glassy carbon working electrode (Antec Leyden) set at +0.7 V with respect to a Ag/AgCl reference electrode. Mobile phase contained 13% MeOH, NaH_2_PO_4_ (0.12 M), EDTA (0.8 mM), OSA (0.5 mM) pH 4.6. The flow rate was fixed at 1⍰mL/min. Analyte measurements were normalized to tissue punch volume (pmol/mm^3^). HPLC data was collected with Clarity (DataApex).

### GAT-3 IHC

Pairs of *SNCA*-OVX mice treated with either saline or PGB (subcutaneously twice daily for 3 days) terminally aneasthetised with an overdose of pentobarbital and transcardially perfused with 20–50⍰mL of phosphate-buffered saline (PBS), followed by 30–50⍰mL of 4% paraformaldehyde (PFA) in 0.1⍰M phosphate buffer, pH 7.4. Brains were removed and stored overnight in 4%PFA, and then coronal sections (50 μm) were cut using a vibrating microtome (Leica VT100S). Sections were stored in PBS with 0.05% sodium azide. Sections were washed in PBS x3 and blocked for 1hr in PBS TritonX (0.3%) with sodium azide (0.02%; PBS-Tx) containing 10% normal donkey serum (NDS). Sections were then incubated in primary antibodies overnight in PBS-Tx with 2% NDS at 4⍰°C. Primary antibodies: rabbit anti-GAT-3 (1:250, Millipore/Chemicon, AB1574). Sections were then incubated in secondary antibodies overnight in PBS-Tx at room temperature (Donkey anti-Rabbit AlexaFluor 488, 1:1000, Invitrogen, A21206. Sections were washed in PBS and then mounted on glass slides and cover-slipped using Vectashield (Vector Labs). Coverslips were sealed using nail varnish and stored at 4⍰°C. Then mounted sections were imaged with an Olympus BX41 microscope with Olympus UC30 camera and filters for appropriate excitation and emission wave lengths (Olympus Medical).

### Behaviour (OFT/PAS/DLC)

Prior to experiments mice were acclimatised to handling and to the experimental room for 3 days prior to experiment. On day of experiment mice were transferred to experimental cages (20 X 41 cm) for 30 minutes. The movements of the mice were tracked using two methods: Photobeam activity system (PAS)-Open field (Bilaney); and video recordings from above using RevotechI1706-P webcam (10fps), video footage was analysed using DeepLabCut pose estimation software, which extracted skeletal mouse representations.

Beam-breaks were binned for 5 seconds (maximum temporal resolution possible) and 30 minutes session was initiated from first beam-break in each chamber. PAS data was extracted from output Microsoft access database and analysed in excel and Graphpad prism. Activity states were determined from the PAS data by plotting the first derivative of the smoothed beam-break vs frame for each animal in Prism (10.2); the number of frames between local minima and maxima of the first derivative were counted and verified as corresponding to activity states by inspecting videoframes of three sessions, blind to analysis and genotype/drug treatment. For corroboration of this analysis please refer to Supplementary data3.

To partition the mouse poses across time into discrete behavioural modules (*syllables*), the Keypoint-Moseq (KPMS) ^64^ algorithm was used on the tracking data generated by DeepLabCut. KMPS is an end-to-end unsupervised method that aims to decouple animal pose information from intrinsic tracking noise using a hierarchical generative probabilistic model. Since KPMS is not currently scale-invariant, tracking data from each recording setup were transformed for downstream analysis. Briefly, the corners of the enclosures in each video setup were used to compute a set of *homography matrices* which apply a linear transformation of the points in frame-coordinates into a shared space across videos.

KPMS follows a two-step process for model fitting which require different sets of user-defined hyperparameters: an initial model fit using an autoregressive Hidden Markov Model, followed by fitting of the full hierarchical model. Default parameters were used for the data extraction and PCA pre-processing steps. Optimal values for the *kappa* hyperparameter (roughly, timescale) were computed by parameter sweep to find a value that produced a median syllable duration of 400-500 ms, which previous work has identified as the most relevant timescale for this sort of discrete behaviours ^64,65^. This resulted in kappa=5x10^-7^. Training of the full model was completed until the median syllable duration stabilised, which in practice was 500 iterations. Syllables with a frequency of greater than 0.5% were used downstream for comparing syllable frequencies across conditions, changepoint analysis, and generation of syllable-syllable transition matrices. Due to body size differences, male and female mice were modelled separately.

## Results

### Gabapentin acutely increases dopamine release in dorsolateral striatum through divergent mechanisms in male versus female mice

We first tested whether or not α_2_δ-ligand gabapentin (GBP) acting in the striatum acutely affected striatal DA release. Incubation of striatal slices with GBP (50 µM) for 30 mins increased extracellular concentration of DA ([DA]_o_) evoked by single pulse electrical stimuli in the dorsolateral striatum (DLS) of both male and female mice, in standard [Ca^2+^]_o_ conditions (2.4 mM) as well as in lower [Ca^2+^]_o_ (1.2 mM). (**Figure 1A,B**, three-way ANOVA: effect of GBP F_1,88_ =9.75 *P=0*.*0024*; Sex F_1,88_ =0.69 *P=0*.*41*; [Ca^2+^]_o_ F_1,88_ =45.13 P<0.0001, interaction GBP*Sex*[Ca^2+^]_o_ F_1,88_ =0.158 *P=0*.*69*). The observed increases in [DA]_o_ was not due to any discernible decrease in DA uptake rates (**Figure 1C**, comparison of *K* of single-phase exponential decay curves: two-way ANOVA: Main effect of GBP F_1,92_=0.055 P=0.82), indicating rather an underlying increase in DA release. Gabapentinoids can limit the ability of some neurons to follow high frequency stimulation ^18^, but GBP did not limit the relationship here between [DA]_o_ and stimulus frequencies of 5-100 Hz during 5-pulse trains (**Figure 1D**, three-way ANOVA: main effect of GBP F_1,6_=0.49 *P=0*.*51*; GBP*Sex*frequency interaction F_1,6_=2.02 *P=0*.*20*, 1.2 mM Ca^2+^).

**Figure 1:**
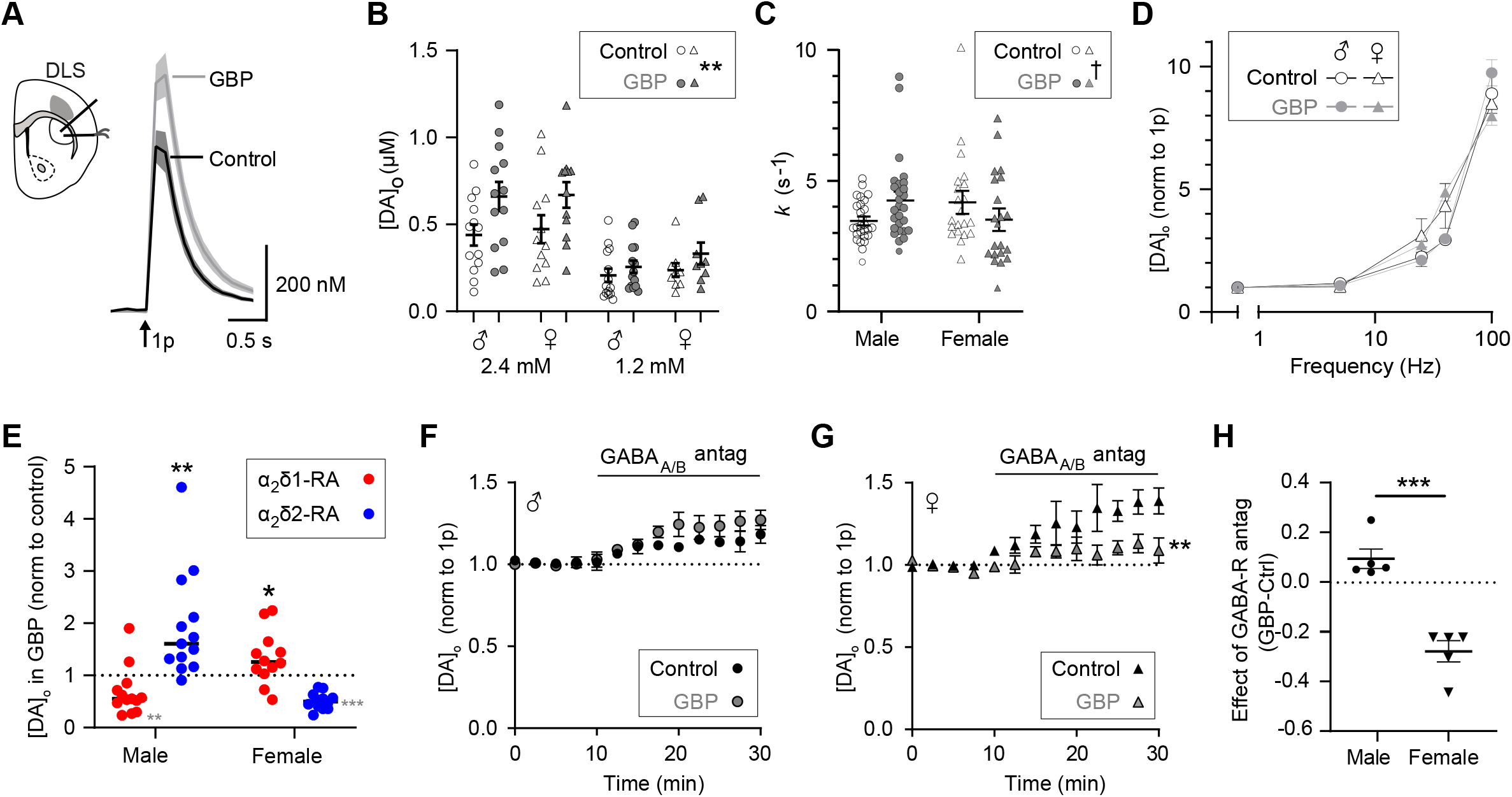
Acute gabapentin increases dopamine release in DLS by divergent mechanisms in male versus female mice. All data was collected in the presence of nAChR antagonist DHβE (1 μM) **(A)** Mean [DA]_o_ ± SEM versus time evoked by single pulses (1p, arrow) in DLS of male and female mice in control (black) and GBP (50 μM, grey) in 2.4 mM [Ca^2+^]_o_ n=25 transients from 10 mice (6m 4 f). **(B)** peak [DA]_o_ in control vs GBP from DLS at 2.4 mM [Ca^2+^]_o_ and 1.2 mM [Ca^2+^]_o_ from male (circle) and female (triangle) mice. Data are per site from male 2.4 N=6, male 1.2 N=4 Female 2.4 N=4 female 1.2 N=3. **(C)** *k* (s^-1^) from single phase exponential decay curves from fastest falling phase from male and female mice in control (black) vs GBP (grey) n=27 female n=21.from both 1.2 and 2.4 mM [Ca^2+^]_o_. ‡ GBPxSex interaction *P=0*.*018* **(D)** mean [DA]_o_ ± SEM normalised to condition 1p vs frequency (for 5p) of male (circle) and female (triangle) mice in control (black) and GBP (grey) conditions. **(E)** Mean [DA]_o_ normalised to control from male and female α_2_δ1-RA (red) and α_2_δ2-RA (blue) mice. Black **(*) significant increase from 1, **(*) significant decrease from 1. **(F**,**G)** [DA]_o_ ± SEM normalised to control vs time (min) in control (black) and GBP (unfilled). GABA_A/B_-antagonists applied at 10 min in DLS of male **(F)** and female **(G)** N=5. **(H)** Effect of GABA-antagonists in GBP subtracted from the effect in control conditions for male (circle) and female (triangle) N=5.

To confirm that the enhancement of evoked [DA]_o_ by GBP involved action at α_2_δ subunits, we utilised α_2_δ mutant mice expressing disrupted RRR motifs that prevent GBP binding, whereby the third arginine residue is mutated to an alanine (R-A mutation), in either the α_2_δ-1 or α_2_δ-2 subunit (respectively α_2_δ1-RA and α_2_δ2-RA mutant mice) ^19,20^. The ability of GBP to enhance evoked [DA]_o_ was prevented in subtype-specific α_2_δ mutant mice in a sex-specific manner; the effect was prevented in α_2_δ1-RA male mice, and in α_2_δ2-RA female mice (**Figure 1E** two-way ANOVA: Sex X Genotype interaction F_1,46_=31.8 P<0.0001). These data confirm that the mechanisms through which GBP promotes DA release require α_2_δ subunits, but α_2_δ1 subtype in male mice and α_2_δ2 in females.

Gabapentinoids typically decrease release of neurotransmitters, by decreasing VGCC density by disruption of forward trafficking of VGCCs into axonal membranes ^21–25^. Since DA release is under a tonic inhibition by striatal GABA release ^26^, we addressed whether GBP enhanced DA release indirectly through a decrease of GABAergic inhibition, a disinhibition mechanism. We assessed the level of GABAergic inhibition of DA release by testing the effects of GABA receptor (GABA-R) antagonists. In male mice, the presence of GBP did not modify the effects of GABA-R antagonists (10 μM bicuculline, 4 μM CGP-55845 on DA release, which increased evoked [DA]_o_ as reported previously by ∼20% (**Figure 1F**, two-way ANOVA interaction GABA-R x GBP F_12,72_=1.22 *P=0*.*28*). By contrast, in female mice, the presence of GBP significantly attenuated the effect of GABA-R antagonists on evoked [DA]_o_ (**Figure 1G**, two way ANOVA F_12,98_=2.91 *P=0*.*0018*) therefore implicating disinhibition as a contributing mechanism through which GBPs increase DA release in female but not male mice. The effects of GABA-R antagonists on evoked [DA]_o_ in the presence versus absence of GBP were significantly different between female and male mice (**Figure 1H**, T-test T_8_=6.4 P=0.0002). These data show that GBP increase evoked DA release in the DLS of male and female mice, albeit via sex specific mechanisms.

### Gabapentin reconfigures VGCCs in male mice

Unlike in female mice, the mechanism by which GBP enhanced DA release in DLS of male mice was not due to disinhibition via GABA-Rs. We therefore sought to explore if GBP is capable of altering the relationship between Ca^2+^ entry and DA release, to identify alternative explanations for the effects of GBP on DA release in male DLS. We next explored whether GBP effects involved changes to how Ca^2+^ and VGCCs supports DA release. We first assessed whether an impact on axonal [Ca^2+^]_i_ could be discerned by imaging GCaMP6f in DLS of DAT-Cre^+/-^:Ai95D^+/-^ mice. This showed that axonal Ca^2+^ transients evoked by a single pulse and their peak dF/F values were unchanged by GBP (50 µM), in male or female mice (**2A**,**B**, two-way ANOVA main effect of GBP F_1,83_=1.67 *P=0*.*20* GBP F_1,83_=0.09 *P=0*.*76*). We then explored in male mice whether GBP modified how VGCCs support DA release. In control conditions, the effects of VGCC-subtype-specific antagonists corroborated that DA release evoked by single pulses in DLS from male mice was supported by N, P/Q, L and T-type VGCCs **(Figure 2C,D)** as seen previously ^5^. Exposure to GBP however, significantly attenuated the effects of PQ-type channel blocker ω-agatoxin IVA (ω-ATX, 200 nM) and L-type inhibitor isradipine (5 µM) on DA release, whilst leaving intact the effects of N-type channel blocker ω-conotoxin GVIA (ω-CTX, 100 nM) or T-type inhibitor NNC 0396 (1 µM) (**Figure 2C,D**, two-way ANOVA interaction VGCC X GBP F_3,27_=4.83 *P=0*.*0081;* effect of GBP F_1,27_=28.24 *P<0*.*0001* post-test isradipine: *P=0*.*0083* ATX: *P<0*.*0001* CTX: *P=0*.*99* NNC: P=0.38), indicating a diminution in the contributions of Ca^2+^ entry via PQ and L-type VGCCs to DA release in male mice.

**Figure 2:**
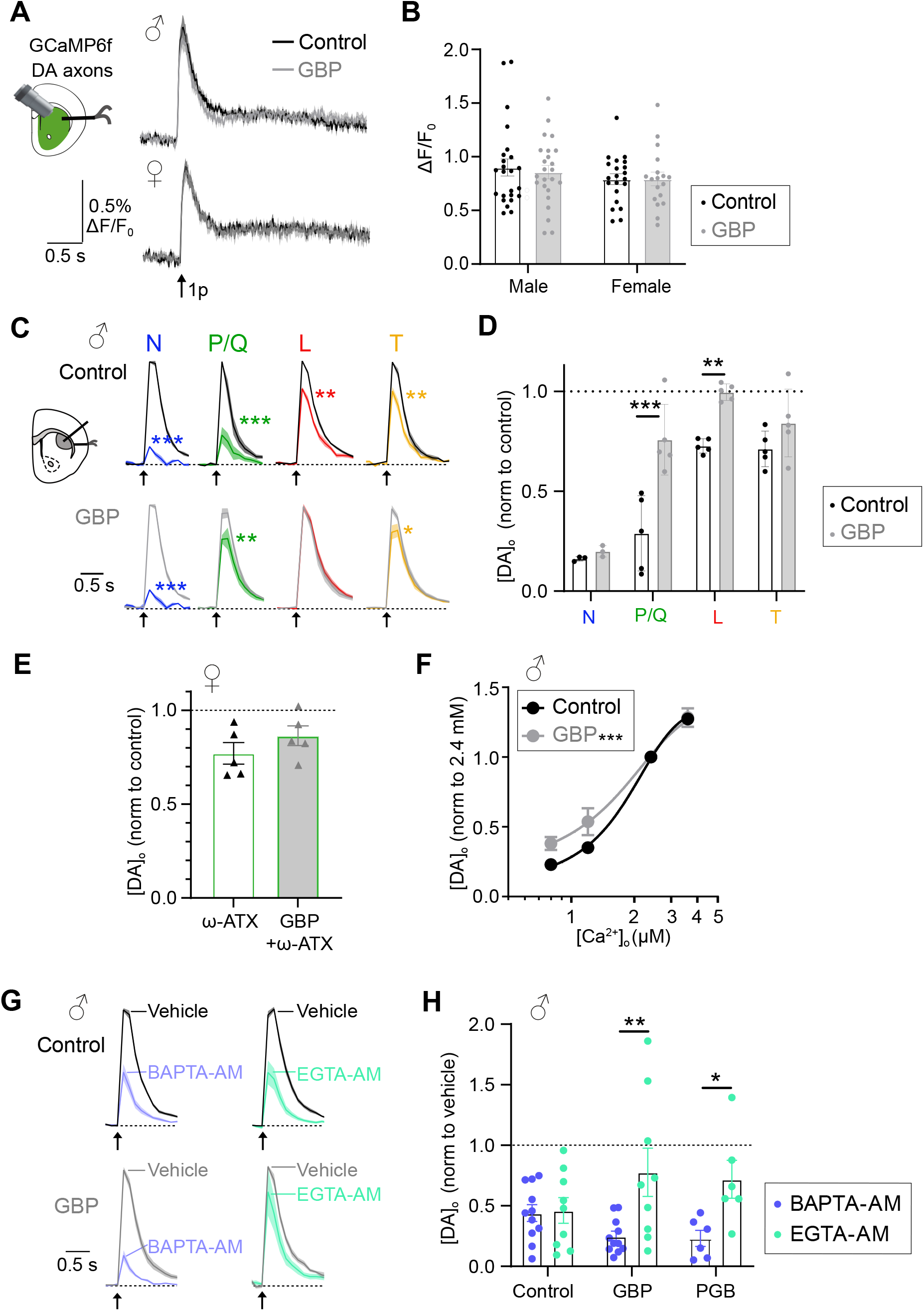
GBP reconfigures spatiotemporal coupling of VGCCs to DA release in male mice. **(A)** Top left: cartoon illustrating recording of GCaMP6f Ca^2+^ signals from male and female DAT-Ccre:Ai95 mice; Ca^2+^ transients recorded in DLS in control (black) and GBP (grey) conditions, n=15 N=5 mice/sex. **(B)** Peak ΔF/F_0_ from male and female mice in control (black) vs GBP (grey). **(C)** Cartoon illustrating FCV recording of DA transients from male WT mice. Mean profiles of [DA]_o_ ± SEM (*shading*) evoked by 1p (*arrow*) in control (*top panel*) or presence of GBP (*bottom panel*) before (black/grey for control and GBP respectively) and after VGCC inhibitors. Data are normalised to peak [DA]_o_ in pre-inhibitor conditions (*black/grey*). N-block, ω-CTX GVIA 100 nM (*blue*); P/Q-inhibitor, ω-ATX IVA 200nM (green); L-inhibitor, isradipine 5 μM (*red*); T-inhibitor block, NNC55-0396 1 μM (*orange*). **(D)** Peak [DA]_o_ evoked in presence of each VGCC inhibitor (N: ω-CTX-GVIA 100 nM (blue); PQ: ω-ATC-IVA 200 nM (blue); L: isradipine 5 µM red; T: NNC-550396 1 µM (orange) without GBP(unfilled) or in presence of GBP (filled bars) normalised to control in absence of VGCC inhibition. **(E)** Peak [DA]_o_ evoked in presence of ω-ATX-IVA from female mice without GBP (unfilled) and or in presence of GBP (filled bars), N=5. **(F)** Peak [DA]_o_ ± SEM evoked by 1p normalised to 2.4 mM [Ca^2+^]_o_ vs [Ca^2+^]_o_ in control (black) and GBP (grey) in DLS of male mice. N=4. (G) Mean profiles of [DA] ± SEM (shading) vs time evoked by 1p in control (upper panel) or presence of GBP (lower panel), from vehicle incubated or either the fast Ca^2+^ chelators BAPTA-AM (*left, blue*) or the slow chelator EGTA-AM (right, green). Data are normalised to vehicle conditions. **(H)** Peak [DA]_o_ release (% of vehicle) following incubation with BAPTA-AM (100 μM) (blues) or EGTA-AM (100 μM) (greens) in control versus GBP. N=4 animals. Two-way ANOVA, Sidak post-tests, * P<0.05, **P<0.01.

We considered comparative impact in female mice. We did not characterise GBP impact on L-type VGCC function in female mice because L-type VGCCs do not support DA release in female mice ^27^. We found modest P/Q channel function in supporting evoked [DA]_o_ in females in control conditions (**Figure 2E**, 29% inhibition by ω-ATX-IVA versus 77% in males, non-contemporaneous and so not statistically compared) but this was not modified by GBP (**Figure 2E** unpaired T-test T_8_=1.20 P=0.26) further supporting a distinction in VGCC function and mode of action of gabapentinoids between sexes. The finding that GBP does not change the effect of ω-ATX-IVA in female mice also excludes an alternative explanation that GBP changes ω-ATX-IVA in male mice through a drug-drug interaction rather than changing P/Q function. Similarly, the effects of isradipine seen in the DLS of male α_2_δ-RA mutants were not occluded by GBP (**Supplementary Figure S1**), supporting our interpretation that GBP is acting through α_2_δ subunits to reconfigure the roles of VGCCs in the control of DA release.

To understand how GBP increases the levels of evoked [DA]_o_ without modifying bulk axonal Ca^2+^ levels but while reducing the contributing roles of some VGCC subtypes in male mice, we tested whether GBP changes the relationship between extracellular [Ca^2+^] ([Ca^2+^]_o_) and DA release. We found that GBP decreased the Hill slope (0.72 vs 0.55) and EC_50_ (63.25 vs 35.01) of the relationship between [Ca^2+^] and [DA] evoked by a single stimulus pulse, leading to relatively potentiated DA release in low [Ca^2+^]_o_ (**Figure 2F**, comparison of fits sigmoidal dose response curve F_2,27_=6.04 *P=0*.*006*). Together, this set of changes to the relationships between [Ca^2+^]_o_, VGCC function and DA release, but not bulk axonal Ca^2+^ levels, suggests that GBP might be modifying the spatiotemporal organisation of Ca^2+^ entry in relation to DA release.

We examined whether GBP, and an alternative gabapentinoid pregabalin (PGB), alter the spatiotemporal coupling of Ca^2+^ to DA release. We utilised the fast and slow exogenous, intracellular Ca^2+^ buffers BAPTA-AM and EGTA-AM respectively. The relative effects of BAPTA-AM and EGTA-AM, on neurotransmitter release can indicate a dependence on Ca^2+^ sources that are tightly/locally coupled versus loosely/remotely coupled to the release machinery. i.e. when the effect of BAPTA is greater than EGTA this indicates tight spatiotemporal coupling ^28^. In control conditions in male mice, as previously published ^5,29^, both BAPTA-AM and EGTA-AM decreased evoked [DA]_o_, with EGTA-AM action indicating relatively loose spatiotemporal coupling between the sources of axonal Ca^2+^ (VGCCs) and the DA release machinery. Incubation with either GBP or PGB significantly increased the relative effect of BAPTA-AM vs EGTA-AM (**Figure 2G,H**, two-way ANOVA interaction: chelator X GBP F_2,46_=3.25 *P=0*.*048* post test control *P=0*.*99* GBP: *P=0*.*0032* PGB: *P=0*.*047*) indicating a tightening of the spatiotemporal coupling between sources of Ca^2+^ to DA release. Together, these findings suggest that in male mice, gabapentinoids act via α_2_δ subunits in DA axons to limit recruitment of PQ- and L-type VGCCs, that these VGCC subtypes are contributing sources of Ca^2+^ that are loosely coupled spatiotemporally to DA release, and that a reconfiguration of VGCCs results in tight and efficient support of DA release by Ca^2+^ without compromising levels of net Ca^2+^ entry or DA release.

### Pregabalin increases dopamine release in DLS of *SNCA*-OVX mice

Our findings show that acute application of GBP to striatum increases DA release in DLS of both male and female mice but through sex-specific mechanisms. In males, GBP reconfigured VGCCs to diminish dependence on L- and P/Q VGCCs but increased efficiency of coupling of Ca^2+^ entry to DA release, while in females, GBP limited an inhibitory GABAergic tone. Despite diversity in mechanisms, the increase in DA release was common to both sexes, and could confer opportunity for gabapentinoids to be of therapeutic benefit to both sexes in the treatment of Parkinson’s.

To better understand this translational value, we addressed whether systemic *in vivo* gabapentinoid dosing promotes striatal DA release to rescue release deficits in a mouse model of PD. Human wild-type α-synuclein-overexpressing mice (SNCA-OVX) ^30^ have a 30% deficit in DA release restricted to DLS, when compared to α-synuclein-null background control mice, from 3 - 20 months of age ^30–32^. We systemically administered PGB rather than GBP owing to the better bioavailability of PGB after systemic dosing ^33^ and used a dose that we estimate approximates to 300 mg/day in patients. PGB (30 mg/kg, s.c.) administered systemically to 12-15-month-old male *SNCA*-OVX mice twice daily for 3 days increased evoked [DA]_o_ detected *ex vivo* in striatal slices by ∼40% in the DLS but not in the ventral striatum, compared to [DA]_o_ evoked in saline vehicle-treated controls (**Figure 3A**). A single *in vivo* dose of PGB administered 1 hour prior to slice preparation was insufficient for this effect (**Figure 3B**, One-way ANOVA F_2,52_=5.86 *P=0*.*005* post-test PGBx6 vs saline *p=0*.*024* PGBx1 vs Saline *P=0*.*653*). We replicated the 3 days of twice daily injections with PGB across broader age ranges (from 3 to 22 months) and for cohorts of all male, all female or mixed sex mice, and found in each case that evoked [DA]_o_ in DLS was increased (by 35-46% depending on cohort) versus vehicle-injected controls (**Figure 3C**, 2-way ANOVA main effect of PGB F_1,206_=31.95 *P<0*.*0001*; interaction cohort x PGB F_3,206_=0.57 *P=0*.*63*). The level of increase in evoked [DA]_o_ equates to a reversal of the underlying DA release deficit.

**Figure 3:**
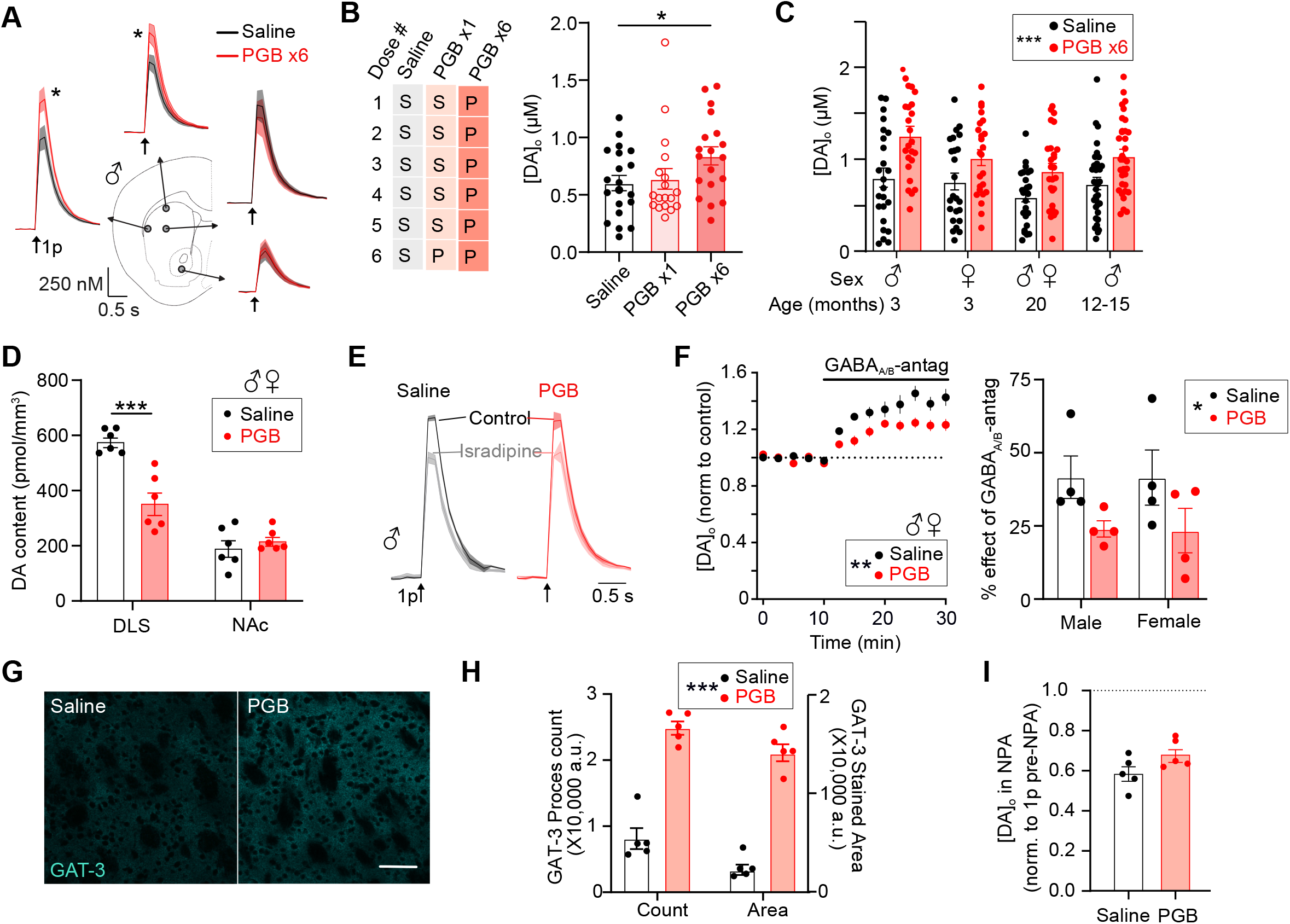
Chronic PGB increases DA release in DLS of _*SNCA*_-OVX mice. **(A)** Evoked DA transients from male 12-15 month old *SNCA*-OVX mice injected with saline vehicle (black) vs PGB (red) (30 mg/kg, 3 days b.d. = 6 doses) from sites across the striatum. N=8 pairs, 16 sites/region. **(B)** Left, dosing regimens; Right, Peak [DA]_o_ evoked by single pulses in DLS from mice injected with saline, PGB x1 doseor PGB x6 doses. N=3 triplicates 18 sites/treatment. **(C)** Peak evoked [DA]_o_ in DLS from mice injected with saline (black) vs PGB (6 doses) across different treatment cohorts. Cohort size: 3 months, 4 pairs/sex; 20 months, 5 pairs (3f, 2m); 12-15 months males, 8 pairs. **(D)** DA content (pmol/mm^3^) of DLS and NAc of *SNCA*-OVX mice injected with saline (black) vs PGB (red). N=6. **(E)** Mean 1p-evoked [DA]_o_ ± SEM vs time (s) from male *SNCA*-OVX mice injected with saline (black) vs PGB (red) in control (bold line) and LTCC-inhibitor isradipine (fine line), normalised to pre-isradipine control levels, N=5. **(F)**(*left*) Mean peak [DA]_o_ ± SEM vs time (min) in DLS of *SNCA*-OVX mice injected with saline (black) vs PGB (red). GABA-R antagonists applied at 10 min, data normalised to pre-antagonist control level. N=7 pairs. (*right*) Summary effect of GABA-R antagonists from male and female mice treated with saline (black) or PGB (red) **(G)** Representative images of GAT-3 staining from DLS of mouse injected with saline vehicle (left) vs PGB (right), n=8 (4 mice), scale bar, 100 μm, **(H)** GAT-3 staining process count and stained area quantified. N=5. **(I)** Peak evoked [DA]_o_ in DLS of *SNCA*-OVX mice injected with saline vehicle (black) vs PGB (red), after acute application of GAT inhibitor NPA, normalised to pre-NPA level. N=5.

Furthermore, *SNCA*-OVX mice store elevated levels of DA compared to controls ^30^, which likely reflects a compensatory response to a deficit in DA release. In line with a reversal of the DA release deficit by PGB, we found that it also significantly lowered DA content of DLS, but not NAc (**Figure 3D** 2-way ANOVA interaction, Region X PGB, F_1,20_=20.02, *P=0*.*0002*, post-test: DLS saline vs PGB, *P<0*.*0001;* NAc saline vs PGB, *P=0*.*76*), approximating to a normalisation of DA content alongside DA release.

To probe mechanisms underlying rescue of DA release by chronic PGB dosing, we assessed whether a downregulation of L-type VGCC function seen in male mice in response to acute administration of GBP (see Fig. 2) was also an outcome of the more chronic systemic dosing. However, application of VGCC inhibitor isradipine to slices taken from saline- or PGB-injected male mice similarly decreased evoked [DA]_o_ (**Figure 3E** comparison of isradipine effect T-test T_8_=1.01 P=0.34), indicating that changes to L-type VGCC function do not contribute to the persistent increase in DA release seen, and likely require gabapentinoids to be acutely present.

*SNCA*-OVX mice have been found to display elevated tonic inhibition of DA release by GABA-Rs in DLS, thought to result from a ∼30% decrease in striatal density of GABA-transporters GAT-1 and GAT-3 ^31^. Given also that acute administration of GBP in females at least acutely reduced tonic GABA inhibition on DA release (see Fig. 1F-H), we tested whether chronic *in vivo* PGB dosing decreased tonic GABA inhibition of DA release, and restored GAT function. Using a mixed sex cohort, we found that *in vivo* dosing with PGB decreased the effect of GABA-R antagonists on evoked [DA]_o_ *ex vivo* (**Figure 3F**, two-way ANOVA interaction, GABA-R antagonists X PGB, F_12,182_=2.76 P=0.0018; post-test, *P<0*.*01 from 22*.*5-30 min – RHS panel:* two-way ANOVA main effect of sex F_1,12_=0.002 *P=0*.*96*, PGB F_1,12_=6.19 *P=0*.*029*)), effectively reversing the elevation in GABA tone in both sexes. PGB also increased striatal density of GAT-3 immunoreactivity (**Figure 3G, H** two-way ANOVA, main effect of PGB F_1,8_=107.8 *p<0*.*001* N=3 male and 2 female), reversing the lowering of GAT levels in this model, and consistent with GAT levels acting to limit GABA tone at its receptors. Intriguingly however, the effect on evoked [DA]_o_ of an inhibitor of GATs (broad-spectrum inhibitor nipecotic acid, NPA) was not enhanced in PGB-compared to saline-treated mice (**Figure 3I** T-test T_8_=2.0 *P=0*.*081*), suggesting that upregulation of GAT-3 levels by PGB might not be the primary mechanisms through which GABA tone is downregulated.

These findings do however suggest a phenotypic convergence of mechanisms in males and females through which a short course of chronic dosing with PGB corrects a DA release deficit in the dorsal striatum and normalise elevated DA content as well as elevated GABAergic inhibition. We note that although this short dosing regimens of PGB corrected these DA and GABA signalling phenotypes of the *SNCA*-OVX mouse, it did not reverse some other adaptations previously reported for this mouse, namely enhanced DAT function^32^ or α-synuclein oligomer numbers per midbrain DA neuron ^34^ (**Supplementary Figure S2**) on our experimental timescale.

### PGB increases behavioural transitions in *SNCA*-OVX mice

We tested whether PGB dosing resulted in improvements to motor function in the *SNCA*-OVX mouse model of PD. We first established an assay that revealed early movement deficits in SNCA-OVX mice of 6 months of age that recapitulate earliest motor symptoms reported in human PD, before testing effects of PGB. At 6 months of age, *SNCA*-OVX mice have a 30% DA release deficit prior to detectable DA neuron cell loss but no motor deficits have been detected previously in test of gross motor function such as latencies to fall from rotarod, or total distance moved in open field test until old age ^30^. Some of the earliest motor symptoms reported in PD are difficulties with initiating movements and with changing speed and direction, consistent with suggested roles for DA in facilitating moment-moment action selection, and having a role in “chunking” behavioural episodes ^35,36^. We revisited motor function in *SNCA*-OVX mice at 6 months of age using a more sensitive photobeam activity system (PAS) and DeepLabCut to analyse moment-by-moment changes to naturalistic movement in a novel environment.

We found that the total number of beam-breaks (indicative of distance travelled) did not differ between the *SNCA*-OVX and *Snca*-null background control mice (**Figure 4A**, two-way ANOVA, Genotype X sex interaction F_1,24_= 0.05 *P=0*.*83*, effect of Genotype F_1,24_=0.83 *P=0*.*37*), but the distribution of the number of beam breaks per 5s significantly differed (**Figure 4B**, Kolmogorov-Smirnov test KS=0.16 *P=0*.*0002*), suggestive of a disturbance in the pattern of movement. To characterise this disturbance, we plotted rates of change in movements (first derivative of smoothed beam-break data) over time (**Figure 4C**) and quantified time between changes in activity states (time interval between each local maxima and minima). We identified that *SNCA*-OVX mice have significantly fewer transitions between activity states (**Figure 4D**, mean number transitions: SNCA-OVX 15.4 ± 0.9, *Snca*-null 19 ± 0.6, T_26_=3.48 *P=0*.*0018*) and greater interval times between states than controls (**Figure 4E**, mean duration: *SNCA*-OVX 99 s, *Snca*-null 79 s, Mann Whitney test, U=22359, *P<0*.*0001*). This indicates that once moving, *SNCA*-OVX mice exhibit fewer changes in their rate of movements vs their Snca-null controls, reminiscent of early movement deficits in PD.

**Figure 4:**
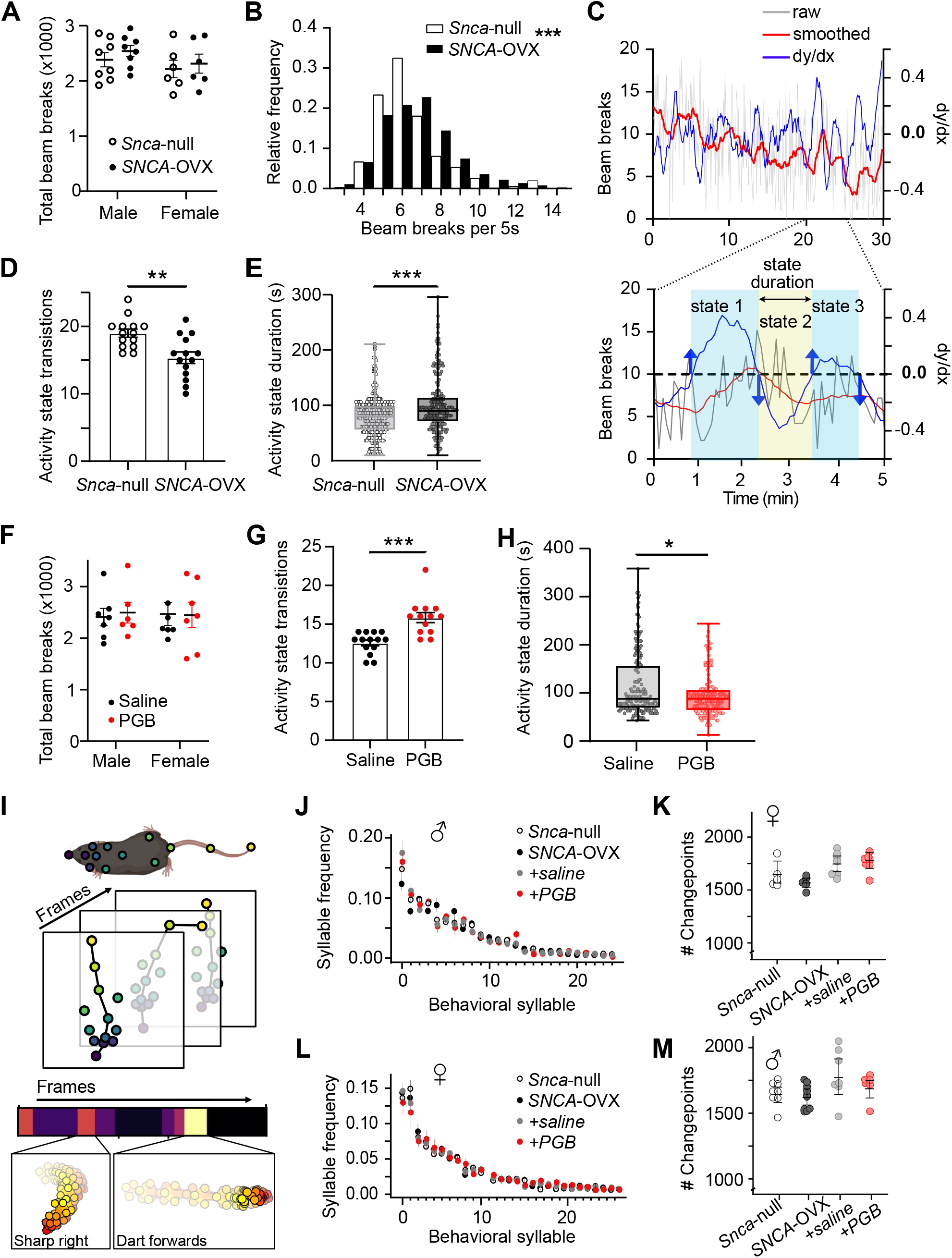
PGB increases behavioural transitions in SNCA-OVX mice. **(A)** Total beam breaks from male and female *Snca*-null (unfilled) vs *SNCA*-OVX (filled) (6 month) mice across 30 min session mice. N=8 male pairs, 6 female pairs. P=0.37. **(B)** histogram showing distribution of beam breaks per 5 sec from *Snca*-null and *SNCA*-OVX mice. **(C)** *upper*: An example 30-minute session, indicating how transition states were calculated. Number of beam breaks (binned across 5 sec) vs frames (5s bins) (grey), smoothed beam break line (red), the 1^st^ derivative (dy/dx) of the smoothed line. *Lower: zoomed in 5 minute window*. Dashed line indicates dy/dx = 0. Upwards blue arrow indicates point when dy/dx crosses from negative to positive and downward blue arrow indicates when dy/dx crosses from positive to negative indicating changes in activity sates. Shaded boxes indicate three designated “activity states” and horizontal arrow indicates sate duration. **(D)** summary of total number of intervals calculated per mouse. N=14. **(E)** length of each interval calculated for mice of each genotype n=268 vs 215 **(F)** Total beam breaks from male and female *SNCA*-OVX mice treated with saline (black) vs PGB (red) 30 mg/kg 3d bd mice across 30 min session. N=6 male pairs, 7 female pairs. P = 0.87. **(G)** summary of total number of intervals calculated per mouse. N=14. **(H)** length of each interval calculated for mice of each genotype n=176 vs 206 **(I)** Cartoon indicating behavioural syllable designation based on DLC bodypoint tracking and KPMS **(J**,**L)** frequency of behavioural syllables for male (J) and female (L) mice **(K**,**M)** number of changepoints identified by KPMS from male (K) and female (M) mice.

We assessed whether PGB (30 mg/kg s.c. twice daily for 3 days) rescued the behavioural phenotypes we observed in *SNCA*-OVX mice (age 6 months). We first tested the total number of beam breaks, and found that PGB did not alter the total number of beam breaks (**Figure 4F**, Two-way ANOVA main effect of treatment F_1,23_=0.026 *P=0*.*87*), but did significantly increase the number of transitions (**Figure 4G**, T_25_=4.45 *P=0*.*0002*) and potentiated shorter state durations (**Figure 4H**, Mann Whitney test U=11695 *P=0*.*010*). We replicated these analyses using DLC body tracking data and found comparable effects in both cohorts (*SNCA*-OVX vs *snca*-null and Saline vs PGB) (**Supplementary Figure S3**)

To interrogate whether the difference in motor patterns were due to changes in subsecond specific behaviours (turns, accelerations, perimeter running etc), we used Keypoint-MoSeq (KPMS) following DeepLabCut body point tracking (**Figure 4I**) to analyse individual behavioural syllables and conduct changepoint frequency analysis to identify moments of sudden, simultaneous changes in the trajectories of multiple bodypoints. We did not however reveal any differences between either genotype or drug condition in frequency of syllables (**Figure 4J,L**) or in the number of changepoints (**Figure 4K,M**) in either male or female mice. These data show that PGB does not perturb normal behaviour patterns, instead improving motor flexibility.

In summary, we identifed a subtle behavioural phenotype in the *SNCA*-OVX mouse model at 6 months of age, prior to neuronal loss, and that an observed impairment to motor flexibility was rescued by treatment with PGB.

## Discussion

Here we have identified that gabapentinoids acutely and chronically regulate striatal DA release in male and female mice, boosting DA release through actions involving VGCC α_2_δ subunits and atypical alterations to the spatiotemporal dynamics of how Ca^2+^ supports DA transmission, as well as disinhibition via reductions to GABA transmission. In turn, we identified an increase in DA transmission and rescue of motor impairments in a mouse model of PD after systemic PGB. These findings establish gabapentinoids as bearing exciting potential utility for repurposing to improve motor function in Parkinson’s, now warranting further study.

### Multiple mechanisms and α_2_δ subtypes

The ability of GBP to increase striatal DA release seems surprising given previous observations that gabapentinoids typically decrease the presynaptic density of VGCCs and limit release probability of other transmitters ^21,24,25^. We reconciled this disparity by identifying two distinct mechanisms through which gabapentinoids increase striatal DA release. Firstly, in male striatum, GBP limited how some VGCC types (L-, P/Q) support DA release but, overall, tightened the spatiotemporal coupling of sources of Ca^2+^ entry to DA release, promoting its efficiency. Secondly, in female striatum acutely exposed to GBP, or when mice of either sex were treated chronically with PGB *in vivo*, there was a disinhibition of DA release due to reduced inhibition by GABA-Rs, consistent with reduced GABA release.

It is unresolved whether these mechanisms are mutually exclusive but convergent, or whether the reconfiguration of VGCCs that occurs when the drug is administered chronically alters the coupling between GABA signalling and DA release. The effects of GBP required sex-specific α_2_δ subunits: α_2_δ1 in male and α_2_δ2 in females. Insufficient comparative expression data exists for males versus females to understand whether this divergence reflects underlying expression. Few studies have investigated the effect of gabapentinoids in female rodents, but one previous study did identify that GBP is less effective at preventing seizures in female than male mouse pups ^38^. Regardless of this divergent dimorphism, ultimately the impact on boosting DA release converged after in vivo dosing through GABA disinhibition. Ultimately, a convergence in outcome provides support for a proposal that gabapentinoids could be considered for repurposing for treating PD in both men and women in order to boost DA function.

Besides setting the level of GABA release, there remain untested possibilities for GBP action to include component mechanisms acting via astrocytes and/or thrombospondins. The level of striatal GABA tone is also set by the level of GABA uptake via GATs, that are at least in part astrocytic^39^, while astrocytes secrete thrombospondins that act at α_2_δ subunits to shape or maintain synapses ^40^, an action that is disrupted by GBP, ^40^ through which gabapentinoids might also stably change the function of DA networks and release.

### Translational potential for Parkinson’s

The increase in DA release by gabapentinoids that we have identified here could have major impact for the symptomatic treatment of Parkinson’s. There is evidence from small scale trials and UPDRS scoring indicating that gabapentinoids can improve symptoms in Parkinson’s, and in parallel, gabapentinoids are a main line of therapy for treating restless-leg syndrome which is associated with DA dysregulation ^15–17,41^. The precise role of DA in the aetiology and treatment of RLS is currently not well understood. Increased DA turnover has been identified in RLS patients, in addition to decreased D2R and membrane-bound DAT levels consistent with decreased fDOPA uptake, which can be interpreted as indicating a hyperdopaminergic state. However, RLS can be treated with DA agonists and L-DOPA, suggesting RLS may be hypodopaminergic disorder^42^. Gabapentinoids are thought to provide pain relief to patients with RLS, however data presented here could indicate that gabapentinoids may prevent an aberrant increased DA synthesis by correcting DA transmission to improve symptoms of RLS. Our data provide a long-awaited explanation for this efficacy: gabapentinoids directly promote striatal dopamine signalling, and thus efforts to repurpose gabapentinoids for PD should be actively reignited without further delay.

To support this further, we found that gabapentinoids increase DA release in a mouse model of early Parkinson’s, and selectively so in the dorsolateral striatum, effectively reversing their DA release deficit. The *SNCA*-OVX mouse line is a highly physiological model of Parkinson’s that recapitulates the genetic and proteinaceous burden of modest α-synuclein overexpression, and presents with a subclinical DA release deficit restricted to dorsal striatum before 3 months of age, and shows gross motor deficits and DA neuron loss in old age ^30,34^ . We found that PGB exposure reversed DA release deficits across all ages, and also normalised associated disturbances in DA content, GABAergic inhibition, and striatal GAT density. We note that while the normalisation of GAT density is consistent with restoration of GABA tone, intriguingly we found that the impact of GAT inhibitors on DA release was not also enhanced suggesting that upregulation of GAT-3 levels by PGB might not be the primary determinants of restored GABA tone. Rather, changes to GAT might be consequential to changes in DA and/or GABA release as seen in GPe ^43^.

We applied a sensitive motor phenotyping method to this model and identified that by 6 months of age, *SNCA*-OVX mice showed fewer changes in their rates of movement than background controls, akin to more inertia. This observation is consistent with evidence supporting an underlying role for DA in building behavioural sequences ^35,36^, and with descriptions of Parkinsonian motor deficits as deficits in initiating and terminating i.e. changing movements. Moreover, we found that systemically administering PGB to *SNCA*-OVX mice reversed these activity disturbances. These changes to behaviours seen with PGB were not linked to changes in frequencies of any particular behavioural syllables, unlike the reported impact of cocaine for comparison ^37^ which has a profound impact to exacerbate DA signalling ^37,44^. While PGB will be affecting all circuits that express α_2_δ1,2 subunits, its action to rescue modest motor impairments is highly consistent with its modest impact on DA signalling that normalises DA release levels.

Besides offering symptomatic therapy, an ideal treatment for Parkinson’s would also offer neuroprotection. Gabapentinoids might offer neuroprotection through multiple means. Their ability to reduce DA storage might reduce some burden associated with DA oxidative damage. DA is a source of oxidative stress to nigral dopaminergic neurons, due to the metabolic production of dopamine-O-quinone, and a high DAT/VMAT ratio in SNc neurons that promotes free cytosolic levels of DA ^45^. But the impact of gabapentinoids on the underlying spatiotemporal handling of Ca^2+^ might offer a particularly advantageous strategy. The routes and handling of Ca^2+^ entry is of particular pertinence: Ca^2+^ entry into DA neurons via L-type VGCCs is associated with vulnerability to degeneration ^46^. Dihydropyridine inhibitors of L-type VGCC function e.g. isradipine have consequently been tested for neuroprotection in clinical trial but without success. But their limited efficacy has several likely explanations, including: patient stages too advanced Parkinson’s to assess neuroprotection; limited brain target access/binding of isradipine with dosing regimens used ^47,48^; and, potential deleterious impact of L-type VGCC inhibition on levels of DA release and/or on supporting Ca^2+^-catalysed mitochondrial ATP generation ^3,27^ offsetting any potential neuroprotective advantage. Gabapentinoids might offer an alternative strategy to reduce Ca^2+^ burden without the confounding impacts of direct channel inhibitors. The DA axons innervating dorsolateral striatum that are most vulnerable to parkinsonian degeneration typically utilise a wide repertoire of VGCCs to support DA release, including N-, PQ, L- and T-type VGCCs, that together support loose spatiotemporal coupling between sources of Ca^2+^ and downstream DA release ^5^. We found here that the net tightening of spatiotemporal coupling by GBP was associated with a reduction in contribution of Ca^2+^ entry via PQ- and L-type VGCCs. Our finding that gabapentinoids limit L-type VGCC contribution to DA release in axons by making them obsolete without directly inhibiting them, and without compromising DA release but moreover boosting it, might fruitfully limit cytosolic Ca^2+^ burden to offer a combined symptomatic and neuroprotective benefit without the deleterious impact on DA seen for direct channel inhibitors.

### Concluding Remarks

Gabapentinoids are clinically approved medications for several disorders including neuropathic pain, sleep disturbances, anxiety and restless-leg syndrome, and possess long-term tolerability and safety profiles. They could readily be repositioned for a potential new treatment for Parkinson’s. Our findings show that gabapentinoids limit the types of VGCCs mediating Ca^2+^ entry into DA axons to support DA release, while paradoxically enhancing the level of DA release, and furthermore, that in a humanized mouse model of PD, *in vivo* dosing with gabapentinoids can enhance levels of DA release and reverse a movement deficit. These findings provide compelling mechanistic support for some available evidence that gabapentinoids having direct symptomatic benefit ^15–17^ while furthermore, modifying Ca^2+^-associated mechanisms associated with neuroprotective potential in Parkinson’s. This potentially dual benefit, to improve DA function for symptomatic relief, while limiting Ca^2+^ burden for potential neuroprotective advantage, is now ripe for direct testing and exploitation. We note additionally that the ability of gabapentinoids to also help with pain, anxiety and sleep is likely to help with those commonly reported comorbidities in PD sufferers.

The underlying mechanism of action on DA involves α_2_δ-1 and α_2_δ-2 subunits. The α_2_δ subtype of most relevance to humans should be defined to encourage development of subtype-specific selective drugs that might offer enhanced specificity of target engagement. Sex differences might be particularly pertinent if α_2_δ-1 versus α_2_δ-2 subunits are to become refined targets in the future as sex differences in our data suggests that they might have different roles in men and women. In any event, our data now provide a key biological rationale and critical impetus to encourage a well-designed clinical trial, that accounts for potential sex differences, to assess the exciting potential for gabapentinoids to have symptomatic and neuroprotective benefit in Parkinson’s.

## Acknowledgements

The work was funded by support to SJC from Parkinson’s UK (J-1403; G-1803), Wellcome Trust Collaborative Award 223202/Z/21/Z, The Medical Research Council DTP, the Clarendon Fund and Christ Church College Oxford, and to N.B-V. by Project PID2021-128210OA-I00 funded by MICIU/AEI /10.13039/501100011033 and by FEDER, EU and RYC2021-034659-I funded by MICIU/AEI /10.13039/501100011033, EU NextGenerationEU/PRTR, and by the Ikerbasque Basque Foundation for Science, EU COFUND H2020-MSCA-COFUND-2020-101034228-WOLFRAM2, and Achucarro Basque Center for Neuroscience.

## Author contributions

Conceptualisation: KRB and SJC

Methodology and study design: KRB, NBV, LB, MEV, SJC

Data collection: KRB, NBV, BOC, LD, RA, EFL, BMR

Analysis: KB, ALH, NBV, BOC, LD, RA, EL, BMR

Visualisation: KRB, ALH, SJC

Writing original draft: KRB, SJC

Writing-review and editing: KRB, SJC, MEW, NBV, EFL

Funding acquisition: SJC, MEW

## Ethics statement

All procedures were licensed to be carried out at the University of Oxford under the UK Animals (Scientific Procedures) Act 1986.

## Competing interests

The authors declare no competing interests

## Data availability

Data for all figures and code is available at: **10.5281/zenodo.14620800**

## Supplementary materials

Materials and Methods

Figure S1 to S3

References 65-79

## Notes

https://doi.org/10.5281/zenodo.14620801

### Competing Interest Statement

The authors have declared no competing interest.

