## Supplementary Figure for "Gabapentinoids promote striatal dopamine release and rescue multiple deficits of a mouse model of early Parkinson’s"

### Supplementary data : Gabapentin promotes striatal dopamine release

#### Gabapentin (GBP) does not occlude the effect of isradipine in male $\alpha_2\delta1$-RA or $\alpha_2\delta2$-RA mutant mice

We excluded the possibility that GBP action in males does not change VGCC function but rather interferes with the binding of PQ- and L-type VGCC inhibitors through drug-drug interactions. We note that whereas N-type inhibitor acts by directly blocking the channel pore, PQ-channel inhibitor ω-ATX binds outside of the channel pore while L-type inhibitor isradipine is a negative allosteric modulator that shifts activation kinetics (Bourinet et al., 1999; Rey et al., 2020; Li et al., 2024). We excluded drug interactions in two ways. Firstly, the persistence of ω-ATX-IVA action on evoked $[DA]_o$ in females described (Figure 2E) excludes a simple drug-drug interaction. Secondly, we found the effects of isradipine were preserved when we tested in either $\alpha_2\delta1$-RA and $\alpha_2\delta2$-RA male mutant mice in the presence of GBP (**Figure S1A** two-way ANOVA main effect GBP $F_{1,16}$=0.09 *P=0.77*), excluding a drug-drug interaction but corroborating an action of GBP via $\alpha_2\delta$ subunits in the loss of VGCC function. In agreement with data in figure 1E, there is an interaction between the effect of GBP and genotype, whereby GBP no longer increases $[DA]_o$ in $\alpha_2\delta1$-RA mice, however in all conditions isradipine decreases $[DA]_o$ corroborating that GBP is not simply occluding the effect of isradipine due to drug-drug interactions (**Figure S1B** three way ANOVA, main effect of isradipine $F_{1,16}$=94.34 *P<0.0001* GBPxGenotype interactions $F_{1,16}$=7.058 *P=0.017*). Transients indicate no apparent effect of GBP or isradipine on the kinetics of the dopamine transients in either genotype (**Figure 1SC)**.

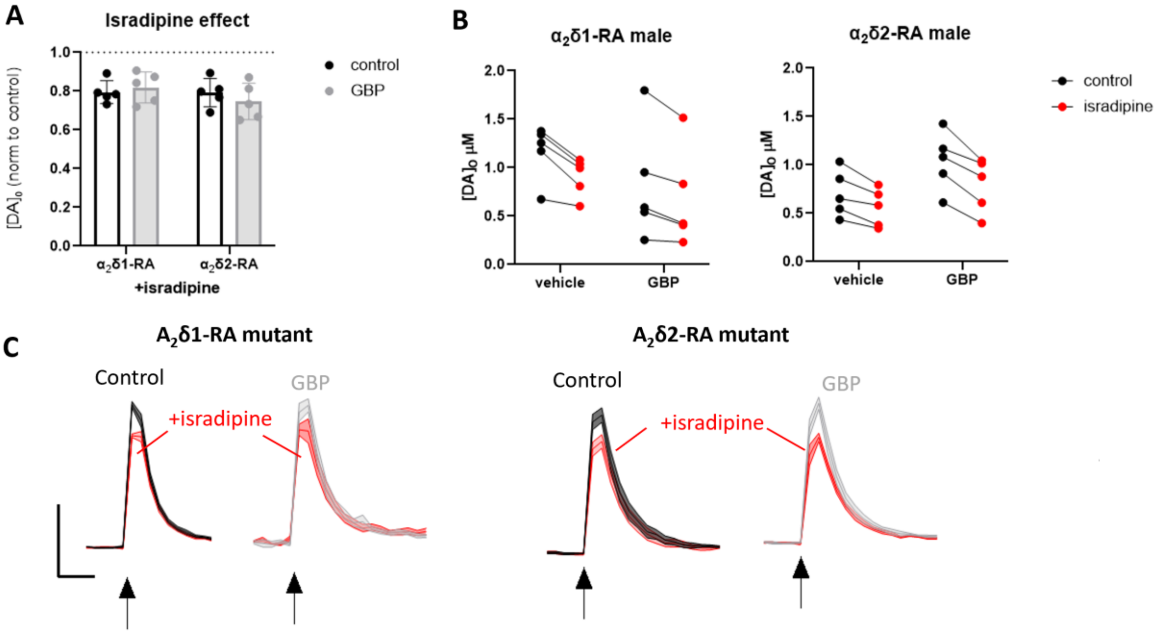


**Figure S1. A)** Summary effect of isradipine in control conditions (black) and following GBP incubation (grey) in the DLS of male $\alpha_2\delta1$-RA and $\alpha_2\delta2$-RA mice. **B)** 1p evoked $[DA]_o$ from $\alpha_2\delta1$-RA (*left*) and $\alpha_2\delta2$-RA mice (*right*) following incubation with vehicle or GBP before and after application of isradipine. **C)** Mean $[DA]_o$ ± SEM versus time evoked by single pulses (1p, arrow) in DLS of male $\alpha_2\delta1$-RA (left) and $\alpha_2\delta2$-RA (right) mice in control (black) and GBP (50 µM, grey) before and after the application of isradipine (5 µM) (red). Scale bar indicates 50% of pre-isradipine control and 0.5s, arrow indicates time of stimulation.

#### Pregabalin does not decrease α-synuclein oligomers or associated changes in DAT function

Despite the role of α-synuclein in the physiology and pathophysiology of DA neurons still being incompletely characterised, α-synuclein is widely assumed to be central to PD-related neurodegeneration. It is becoming increasingly apparent that different molecular forms of α-synuclein can affect different cellular processes and are differently associated with disease pathology. In the *SNCA*-OVX mouse line we identified intracellular α-synuclein oligomers detected with PLA (Bengoa-Vergniory et al., 2020), but could not detect Lewy-like proteinaceous inclusions (Janezic et al., 2013).

It is known that there is a bi-directional relationship between α-synuclein and $Ca^{2+}$, whereby α-synuclein can affect $Ca^{2+}$ signalling including forming $Ca^{2+}$ pores; and $Ca^{2+}$ levels can affect α-synuclein aggregation (Nath et al., 2011; Melachroinou et al., 2013; Ludtmann et al., 2018). Both α-synuclein and $Ca^{2+}$ affect a broad spectrum of DA dependent biology including the DA transporter (DAT) (Lee et al., 2001; Kile et al., 2010; Longhena et al., 2018). We have previously illustrated and interesting link between α-synuclein and DAT function in the *SNCA*-OVX mouse model, whereby α-synuclein promotes cholesterol efflux via ABCA1 transporter, resulting in enhanced DA clearance via DAT, and therefore an enhanced stimulatory effect of cocaine on electrically evoked DA release (Threlfell et al., 2021).

We therefore wanted to test if PGB, by putatively affecting VGCCs function could rescue the aberrant association between α-synuclein and DAT regulation of DA release. We first assessed if PGB administration affected α-synuclein oligomers detectable with a proximity ligation assay (PLA) (Bengoa-Vergniory et al., 2020). A-synuclein-PLA utilises oligonucleotide-conjugated antibodies that, when in close proximity (<16 nm), allow rolling amplification of ligated probes, which can be detected using complimentary fluorescently labelled probes, enabling detection of closely associated antibody targets such as α-synuclein. PGB treatment did not however decrease the number of α-synuclein puncta detected with PLA (**Figure S2A,B** T-test $T_{10}$=1.36 *P=0.20*). Consistent with the finding that PGB did not change α-synuclein oligomers detected with PLA (Figure S2A,B), PGB also did not decrease the effect of cocaine relative to saline-treated animals (**Figure S2C** comparison of cocaine effect T-test $T_4$=0.96 *P=0.39*), indicating that PGB did not change the relationship between DAT and dopamine release. Further experiments are needed to fully determine if PGB is capable of modifying α-synuclein levels, and may require a longer administration period or an alternative method of α-synuclein detection.

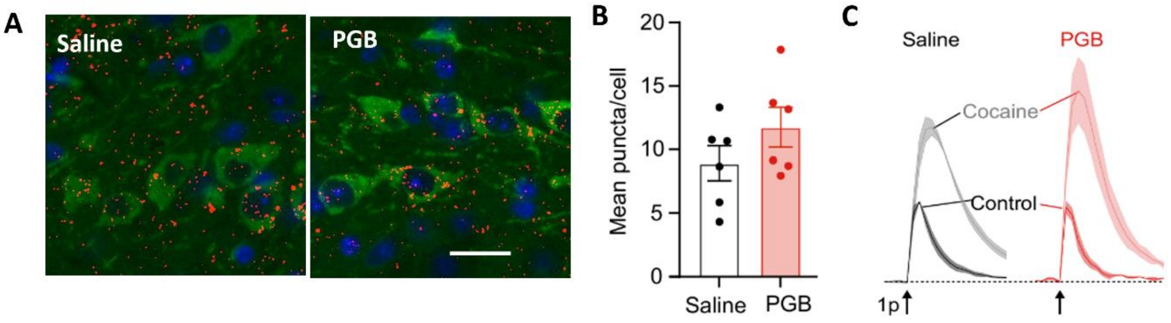


**Figure S2 A:** representative images of α-synuclein PLA puncta (red) per TH+ve cell (green). Scale bar = 50 µm **(B)** PLA signal: mean puncta/cell measured from midbrain of *SNCA*-OVX mice treated with saline vs PGB (6 doses). N=6. **(C)** Mean normalised $[DA]_o$ ± SEM vs time (s) from male *SNCA*-OVX mice treated with saline (black) vs PGB (red) in control (bold line) and DAT-inhibitor cocaine (fine line). N=3.

#### Validation of transition state analysis using DLC data

The relationship between dopamine and motor output is far from trivial. It is well established that the use of dopamine-selective toxins such as 6-OHDA and MPTP leads to striatal dopamine depletion, loss of nigral dopaminergic cells, and profound impairments to locomotion; and conversely use of DA-transporter inhibitors such as cocaine, enhance locomotion. However more recent studies using nuanced genetic approaches have shown that complete ablation of phasic striatal dopamine release does not affect gross locomotion(González-Rodríguez et al., 2021; Delignat-Lavaud et al., 2023; Cai et al., 2024). Moreover, using photometry techniques to detect striatal dopamine release in awake behaving mice has shown a highly nuanced and non-simple relationship between dopamine signalling and motor behaviour (Da Silva et al., 2018; Markowitz et al., 2018; Jørgensen et al., 2023), with some studies indicating roles of dopamine in regulating action output occurring over relatively prolonged time-scales(Howe et al., 2013; Liu et al., 2022).

Here we aimed to test the hypothesis that the DA deficit identified in *SNCA*-OVX mice may result in subtle changes in the behavioural patterns of the mice, in keeping with the observation at early disease stages, people with Parkinson's often report difficulties in initiating or changing movements, describing it as a feeling of "being stuck to the floor". We therefore wanted to detect patterns in motor behaviours of freely moving mice in an open field. In figure 4 we described how using PAS beam-breaks as proxy for distance travelled we could measure differences in the patterns of motor activity, whereby *SNCA*-OVX mice showed fewer transitions between relative activity states, which was rescued by PGB treatment. We replicated the analysis approach using the DLC-tracked data, allowing us to measure actual distance travelled, rather than using beam-breaks as a proxy. Analogously to the approach taken with the PAS beam-break data, we smoothed the raw 10 Hz body tracked data and plotted dy/dx, where a change from a negative value to a positive value, indicated mice had transitioned to an increased activity state and conversely, when the dy/dx went from a positive value to a negative value it indicated a relatively lower activity state **(Figure S3A**). We further corroborated the transitions labelled using this approach, by comparing the time-points identified as changing activity states with a Bayesian changepoint analysis (Rbeast package in R) on the DLC tracked "BodyCentre" point data. The changepoints with the highest probabilities corresponded to the same timepoint identified our transitions data (**Figure S3A** *note that apparent changepoint at ~1450s is not identified by Rbeast*). We chose not to use the Bayesian changepoints for formal analysis because the number of changepoints detected was completely dependent upon the set parameters. And the maximum number of changepoints specified. i.e. if the maximum was set to 20, 20 was found for each session (true if max parameters were set at 10-150), and did not identify some changes that were apparent by eye )e.g. in figure S3A there is an apparent change in activity at ~14375 frames, which is detected by our dy/dx approach, but not identified as a changepoint).

In agreement with data in Figure 4 D-H, we found that *SNCA*-OVX mice have fewer number of transitions (**Figure S3B** Two-way ANOVA effect of genotype $F_{1,22}$=8.16 *P=0.0092*; effect of sex $F_{1,22}$=4.93 *P=0.037*, GenotypeXsex interaction $F_{1,22}$=0.015 *P=0.90***)** , and the average duration of transitions is longer (**Figure S3C** Two-way ANOVA effect of genotype $F_{1,22}$=8.21 *P=0.0090*; effect of sex $F_{1,22}$=4.62 *P=0.043*, GenotypeXsex interaction $F_{1,22}$=0.015 *P=0.91* ). We also found that PGB enhanced the number of transitions (**Figure S3D** Two-way ANOVA effect of treatment $F_{1,20}$=6.03 *P=0.023*; effect of sex $F_{1,20}$=0.97 *P=0.33*, TreatmentXsex interaction $F_{1,20}$=1.72 *P=0.21*) and decreased the average length of transitions (**Figure S3E** Two-way ANOVA effect of treatment $F_{1,20}$=5.97 *P=0.024*; effect of sex $F_{1,20}$=0.53 *P=0.48*, TreatmentXsex interaction $F_{1,20}$=1.53 *P=0.23*)

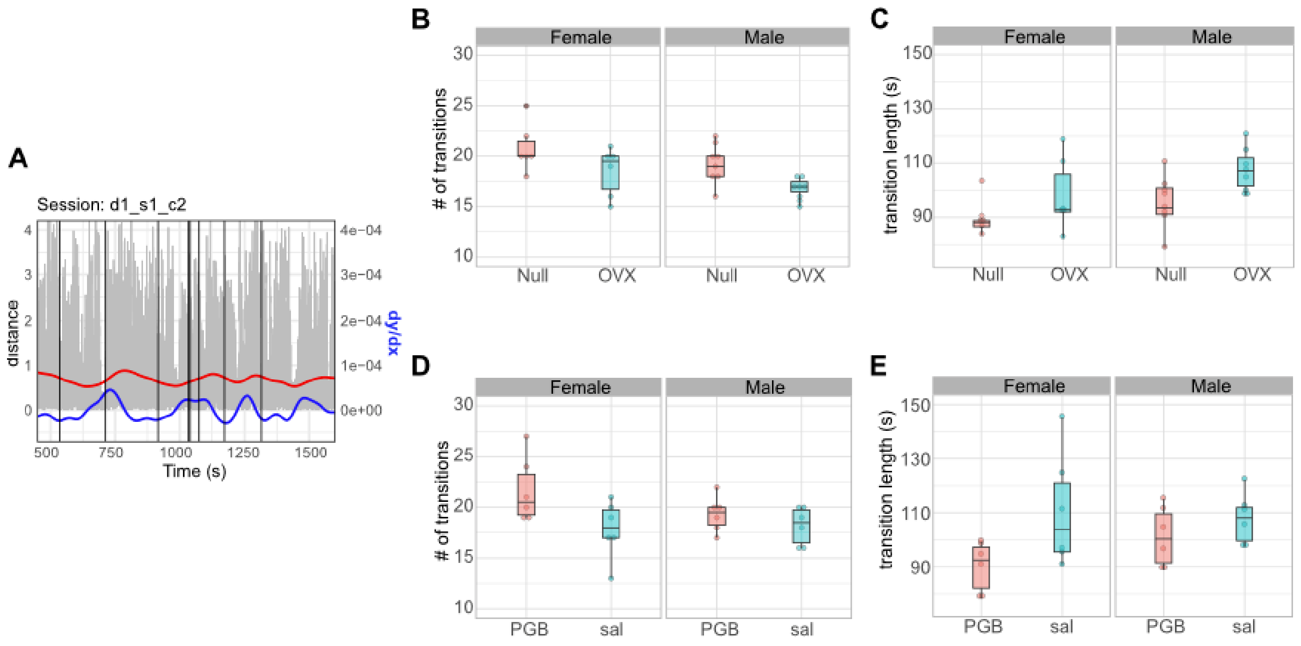


**Figure S3 A:** Representative trace showing raw distance travelled in 10 ms (grey) and the smoothed distance (red) with the dy/dx of the smoothed trace (blue) Vs time (frames every 10 ms). Black vertical lines show changepoints identified with Bayesian change point function Rbeast. **B,D)** summary of number of state transitions identified in each genotype (B) or treatment group (D). **(C,E)** average length of transitions per mouse in each genotype (C) or treatment (E).

#### Supplementary methods

##### PLA

To minimise the number of mice used for the study, following behavioural assays, mice were terminally anaesthetised with pentobarbital, and transcardially perfused with 4% PFA. As previously published (Bengoa-Vergniory et al., 2020) brains were then transferred into 15 ml 4% PFA and stored at 4°C rocking overnight. Brains were then transferred to 70% ethanol at 4°C and left for 24hrs prior to embedding. Paraffin-embedded tissue was dewaxed by 2 min consecutive incubations in Xylene, Histoclear, 100% ethanol, 95% ethanol, 70% ethanol, and H2O. After rehydration, samples were incubated in 10% H2O2 in PBS, to reduce background and heated in a microwave in citrate buffer pH 6.0 (Abcam) for antigen retrieval. After antigen retrieval, samples were subjected to immunofluorescence or AS-PLA as necessary. AS-PLA experiments were performed using the Duolink kits (Sigma) according to the manufacturer's instructions17,18. An α-syn antibody (mouse monoclonal anti-α-syn4D6, ab1903, Abcam, 1:2000), was used to prepare conjugates using Duolink Probemaker Plus and Minus kits. All samples were incubated with Duolink block solution at 37 °C for

1 h and then with α-syn conjugates diluted in Duolink PLA diluent overnight (ON) at 4 °C. Samples were washed with tris buffered saline (TBS) containing 0.05% Tween-20 (TBS-T) and incubated with Duolink ligation reagents for 1 h at 37 °C, washed four times with TBS-T, and then incubated with Duolink amplification reagents for 2.5 h at 37 °C. Samples were washed and then mounted in FluorSave (Calbiochem).

#### Extended behavioural analysis

Experimental protocol is from the main manuscript. DLC body tracking used previously published approach by (Sturman et al., 2020) Sturman et al and "Tracking" analysis pipeline written by Lukaz Von Zeigler. Adapted pipeline to analyse changepoints and transition states were written in R(v4.5.1) (RStudio/2025.05.1+513) all data and code are available at **10.5281/zenodo.14620801.**


Bourinet E, Soong TW, Sutton K, Slaymaker S, Mathews E, Monteil A, Zamponi GW, Nargeot J, Snutch TP (1999) Splicing of α(1A) subunit gene generates phenotypic variants of P- and Q-type calcium channels. Nat Neurosci 2.

Cai X, Liu C, Tsutsui-Kimura I, Lee JH, Guo C, Banerjee A, Lee J, Amo R, Xie Y, Patriarchi T, Li Y, Watabe-Uchida M, Uchida N, Kaeser PS (2024) Dopamine dynamics are dispensable for movement but promote reward responses. Nature.

Delignat-Lavaud B, Kano J, Ducrot C, Massé I, Mukherjee S, Giguère N, Moquin L, Lévesque C, Burke S, Denis R, Bourque MJ, Tchung A, Rosa-Neto P, Lévesque D, De Beaumont L, Trudeau LÉ (2023) Synaptotagmin-1-dependent phasic axonal dopamine release is dispensable for basic motor behaviors in mice. Nat Commun 14.

González-Rodríguez P, Zampese E, Stout KA, Guzman JN, Ilijic E, Yang B, Tkatch T, Stavarache MA, Wokosin DL, Gao L, Kaplitt MG, López-Barneo J, Schumacker PT, Surmeier DJ (2021) Disruption of mitochondrial complex I induces progressive parkinsonism. Nature 599.

Howe MW, Tierney PL, Sandberg SG, Phillips PEM, Graybiel AM (2013) Prolonged dopamine signalling in striatum signals proximity and value of distant rewards. Nature 500.

Kile BM, Guillot TS, Venton BJ, Wetsel WC, Augustine GJ, Wightman RM (2010) Synapsins differentially control dopamine and serotonin release. J Neurosci 30:9762–9770.

Lee FJS, Lie F, Pristupa ZB, Niznik HB (2001) Direct binding and functional coupling of α-synuclein to the dopamine transporters accelerate dopamine-induced apoptosis. FASEB J 15:916–926.

Li Z, Cong Y, Wu T, Wang T, Lou X, Yang X, Yan N (2024) Structural basis for different ω-agatoxin IVA sensitivities of the P-type and Q-type Cav2.1 channels. Cell Res.

Liu H, Melani R, Sankaramanchi A, Zeng R, Maltese M, Martin JR, Tritsch NX (2022) A permissive role for dopamine in the production of vigorous movements. bioRxiv.

Longhena F, Faustini G, Missale C, Pizzi M, Bellucci A (2018) Dopamine transporter/α-synuclein complexes are altered in the post mortem caudate putamen of Parkinson's disease: An in situ proximity ligation assay study. Int J Mol Sci 19.

Ludtmann MHR et al. (2018) α-synuclein oligomers interact with ATP synthase and open the permeability transition pore in Parkinson's disease. Nat Commun 9:2293.

Melachroinou K, Xilouri M, Emmanouilidou E, Masgrau R, Papazafiri P, Stefanis L, Vekrellis K (2013) Deregulation of calcium homeostasis mediates secreted α–synuclein-induced neurotoxicity. Neurobiol Aging 34:2853–2865.

Nath S, Goodwin J, Engelborghs Y, Pountney DL (2011) Raised calcium promotes α-synuclein aggregate formation. Mol Cell Neurosci 46:516–526.

Rey S, Maton G, Satake S, Llano I, Kang S, Surmeier DJ, Silverman RB, Collin T (2020) Physiological involvement of presynaptic L-type voltage-dependent calcium channels in GABA release of cerebellar molecular layer interneurons. J Neurochem 155:390–402.

Sturman O, von Ziegler L, Schläppi C, Akyol F, Privitera M, Slominski D, Grimm C, Thieren L, Zerbi V, Grewe B, Bohacek J (2020) Deep learning-based behavioral analysis reaches human accuracy and is capable of outperforming commercial solutions. Neuropsychopharmacology 45:1942–1952.

Threlfell S, Mohammadi AS, Ryan BJ, Connor-Robson N, Platt NJ, Anand R, Serres F, Sharp T, Bengoa-Vergniory N, Wade-Martins R, Ewing A, Cragg SJ, Brimblecombe KR (2021) Striatal Dopamine Transporter Function Is Facilitated by Converging Biology of α-Synuclein and Cholesterol. Front Cell Neurosci 15.
